# Sustained hyperglycemia specifically targets translation of mRNAs for insulin secretion

**DOI:** 10.1101/2023.09.29.560203

**Authors:** Abigael Cheruiyot, Jennifer Hollister-Lock, Brooke Sullivan, Hui Pan, Jonathan M. Dreyfuss, Susan Bonner-Weir, Jean E. Schaffer

## Abstract

Pancreatic β-cells are specialized for coupling glucose metabolism to insulin peptide production and secretion. Acute glucose exposure robustly and coordinately increases translation of proinsulin and proteins required for secretion of mature insulin peptide. By contrast, chronically elevated glucose levels that occur during diabetes impair β-cell insulin secretion and have been shown experimentally to suppress insulin translation. Whether translation of other genes critical for insulin secretion are similarly downregulated by chronic high glucose is unknown. Here, we used high-throughput ribosome profiling and nascent proteomics in MIN6 insulinoma cells to elucidate the genome-wide impact of sustained high glucose on β-cell mRNA translation. Prior to induction of ER stress or suppression of global translation, sustained high glucose suppressed glucose-stimulated insulin secretion and downregulated translation of not only insulin, but also of mRNAs related to insulin secretory granule formation, exocytosis, and metabolism-coupled insulin secretion. Translation of these mRNAs was also downregulated in primary rat and human islets following *ex-vivo* incubation with sustained high glucose and in an *in vivo* model of chronic mild hyperglycemia. Furthermore, translational downregulation decreased cellular abundance of these proteins. Our findings uncover a translational regulatory circuit during β-cell glucose toxicity that impairs expression of proteins with critical roles in β-cell function.

## Introduction

Diabetes results from pancreatic β-cell failure to secrete sufficient insulin to regulate glucose homeostasis. Progressive decline in β-cell function occurs in the setting of hyperglycemia during the progression from early appearance of autoantibodies to frank type 1 diabetes, and during the evolution from compensated insulin resistance to type 2 diabetes (1, 2). In type 1 diabetes, intensive insulin therapy that restores normal glycemic levels increases stimulated C-peptide levels, a reflection of improved insulin biosynthesis and preserved β-cell function (3). In patients with newly diagnosed type 2 diabetes, intensive short-term insulin therapy improves β-cell function and long-term glycemic control (4, 5). These observations support the hypothesis that glucose toxicity contributes to decline in insulin production in diabetes.

β-cells are specialized to couple glucose metabolism not only to insulin secretion, but also to robust insulin peptide production. In response to an acute physiological rise in glucose, proinsulin mRNA translation rapidly increases within 30 to 60 minutes, without change in insulin mRNA abundance (6). This translational regulation requires sequences predicted to form a stem-loop structure within the 5’-untranslated region of the insulin mRNA that bind to protein factors in a glucose-dependent fashion (7). In addition to insulin, glucose acutely upregulates translation of proteins involved in glucose metabolism, insulin processing, secretory granule biogenesis, and insulin exocytosis, without causing an equivalent increase in total protein synthesis (8–11). Given that newly synthesized insulin is preferentially released initially, this mRNA translational program supports physiological increases in glucose-stimulated insulin secretion (GSIS) following brief exposures to high glucose (12).

By contrast, persistently elevated glucose impairs GSIS and insulin translation. Among obese subjects, GSIS decreases with increasing plasma glucose area under the curve in a 3-hour oral glucose tolerance test, without change in insulin sensitivity (13). *Ex vivo* incubation of isolated human or rodent islets over one week in media containing high glucose also impairs GSIS and inhibits translation of proinsulin (14, 15). The impact of sustained high glucose on GSIS in cadaveric human islets is apparent as early as 2 days following exposure to high glucose, a time point at which impairment is reversible, associated with only modest transcriptomic changes, and without evidence for ER stress (16). Sustained exposure of insulinoma cells and isolated islets to high concentrations of glucose and saturated fatty acids to model the metabolic stress of type 2 diabetes increases translation of JUND, a transcriptional regulator of β-cell apoptosis, and induces ER stress (17, 18). However, it is not known whether sustained high glucose alone initiates programmatic regulation of translation prior to induction of ER stress.

Given the importance of translational regulation in GSIS, we sought to gain insight into the impact of sustained glucose elevation on genome-wide β-cell mRNA translation. Using complementary high-throughput approaches in a MIN6 model and validating our findings in primary isolated islets *ex vivo* and in an *in vivo* model of hyperglycemia, we show that chronic glucose excess coordinately downregulates translation of genes that function in metabolism-coupled insulin secretion.

## Results

### Sustained high glucose impairs insulin translation independent of ER stress

Exposure of primary isolated human or rat islets to high glucose for one week increases basal secretion and impairs glucose-stimulated insulin secretion in human and rodent islets (14, 15, 19). We cultured hand-picked isolated primary rat islets in media containing 16.7 mM glucose versus 5.5 mM glucose. Islets were then rested in media with 2.8 mM glucose for 1 hour prior to assaying for GSIS (Figure 1A). Compared to low glucose, islets incubated in high glucose for only 4 days had increased basal insulin secretion (Figure 1B). Although the amount of insulin secreted following stimulation trended up in islets exposed to sustained high glucose, stimulation index was decreased by 69% (Figure 1B), without impact on total islet insulin content (Figure 1C). Shorter incubations in high glucose did not decrease GSIS (not shown). Exposure to high glucose over 4 days did not cause dedifferentiation or transdifferentiation, as expression of β-cell identity genes was similar between the two glucose conditions (Figure 1D). Although global protein synthesis was unchanged between the glucose conditions (Figure 1E), sustained high glucose decreased the rate of insulin synthesis (Figure 1F). Pancreatic β-cells are particularly susceptible to increased ER stress and prolonged ER stress is detrimental to β-cell function (20, 21). However, treatment with high glucose over 4 days did not increase the phosphorylation of PERK or expression of ATF4, well established upstream regulators of ER stress (Figure 1G). Thus, 4 days exposure to sustained high glucose decreased translation of insulin prior to suppression of global protein synthesis, compromised β-cell identity, or sustained engagement of the unfolded protein response pathway.

**Figure 1.**
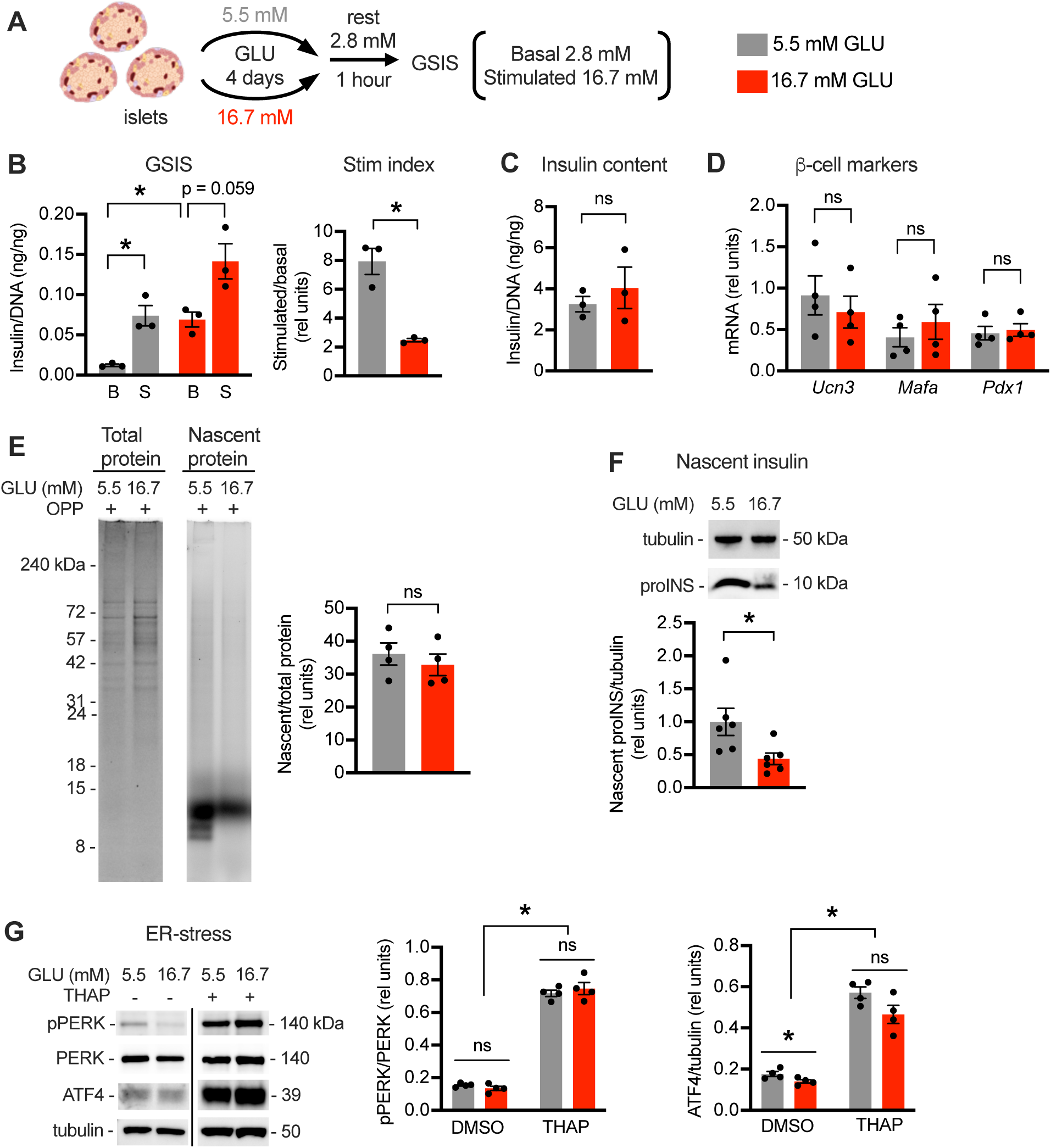
Sustained high glucose decreases basal insulin translation in isolated rat islets. Hand-picked islets from adult Sprague-Dawley rats cultured for 4 days in media containing 5.5 mM (gray) or 16.7 mM (red) glucose (GLU). (**A**) Following 1 hour rest in 2.8 mM GLU, GSIS quantified at 2.8 mM (basal, B) and 16.7 mM (stimulated, S) GLU. (**B**) GSIS normalized by DNA with stimulation (stim) index quantified as stimulatory/basal secretion. (**C**) Insulin content normalized to DNA. (**D**) qPCR quantification of beta-cell markers *Ucn3*, *Mafa*, and *Pdx1* relative to 18S rRNA. (**E**) Cells pulse labeled with O-propargyl-puromycin (OPP) and analyzed by SDS-PAGE. Total protein quantified by Coomassie stain (left) and newly synthesized protein quantified by click-addition of Alexa-647 (right). Representative images with quantification. (**F**) Following metabolic labeling with OPP, click-biotin addition and streptavidin pulldown of nascent proteins. Immunoblot analysis of newly synthesized proinsulin (proINS) with tubulin control. (**G**) Representative immunoblots and quantification for ER stress markers with thapsigargin (THAP)-treated control. Means ± standard error (SE) for n = 3–4 independent experiments. *, P < 0.05 by unpaired or paired t-test. ns, not significant.

### MIN6 cells model chronic high glucose effects on islet insulin translation

Pancreatic islets are micro-organs that consist of several cell types including glucagon-containing α-cells, somatostatin-containing δ-cells, and polypeptide-producing PP-cells in addition to insulin-producing β-cells. To delineate the effects of chronic high glucose specifically on β-cells and to identify a system that more readily provides sufficient material for high-throughput analyses, we modeled chronic high glucose exposure in early passage MIN6 insulinoma cells that support robust GSIS (22). These cells are typically propagated in 25 mM glucose to sustain rapid cell growth but can be maintained for limited periods in lower glucose with slower growth. Following incubation for 24 hours in 5.5 mM glucose, stimulatory glucose caused a 10-fold increase in insulin secretion (Figure 2A-B). Basal insulin secretion did not increase in MIN6 cells maintained in high glucose contrary to observations in islets. However similar to islets, the stimulation index was decreased by 59% by high glucose (Figure 2B). Although high glucose decreased insulin content of MIN6 cells, the impact of sustained high glucose on GSIS remained significant even when secretion was calculated as a percent of insulin content (Figure 2C-D). This did not reflect a general response to increased osmolarity, since incubation in low glucose media supplemented with mannitol did not recapitulate the effect of sustained high glucose (Figure 2E). As observed in rat islets, β-cell identity markers were similar in both glucose conditions, and sustained high glucose specifically decreased insulin translation without affecting global protein synthesis or inducing ER stress (Figure 2F-I). Taken together, these results demonstrate that 24-hour treatment of MIN6 cells with 25 mM glucose vs 5.5 mM glucose models the effects of sustained high glucose on GSIS and insulin mRNA translation in primary islets.

**Figure 2.**
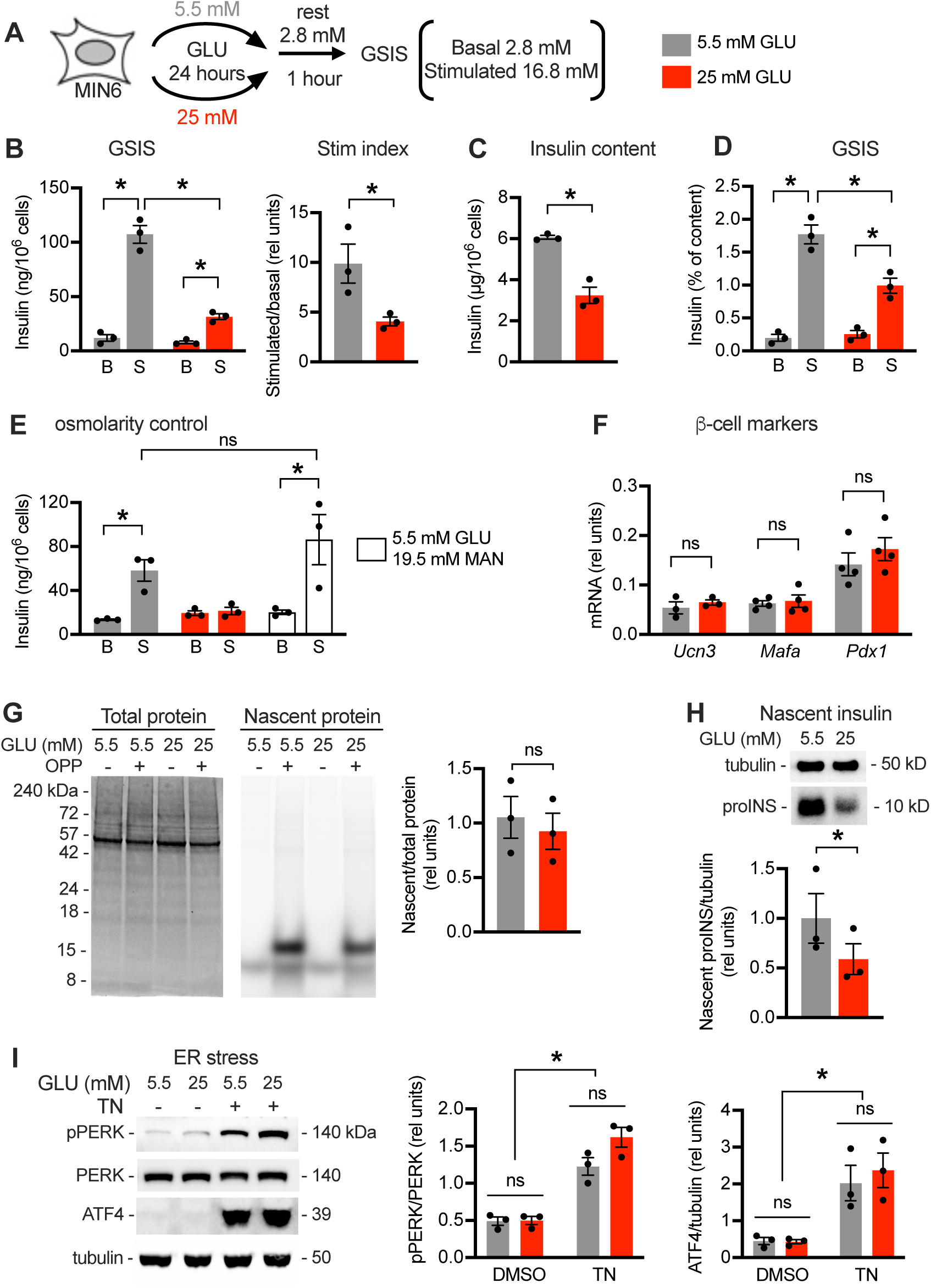
Sustained high glucose decreases insulin synthesis in MIN6 cells. MIN6 cells incubated in media containing 5.5 mM (gray) or 25 mM (red) GLU for 24 hours. (**A**) Following 1 hour rest in 2.8 mM GLU, GSIS quantified at 2.8 mM (B) and 16.8 mM (S) GLU. (**B**) GSIS normalized by cell number with stim index quantified as stimulatory/basal secretion. (**C**) Insulin content per 10^6^ cells. (**D**) GSIS normalized to cellular insulin content. (**E**) GSIS normalized by cell number following incubation in 5.5 mM GLU, 25 mM GLU, or 5.5 mM GLU with 19.5 mM mannitol (MAN, open bars). (**F**) qPCR quantification of beta-cell markers *Ucn3*, *Mafa*, and *Pdx1* relative to 18S rRNA. (**G**) OPP pulse labeling and SDS-PAGE. Total protein quantified by Coomassie stain and newly synthesized protein quantified by click-addition of Alexa-647. Representative images with quantification. (**H**) OPP labeling, click-biotin addition, and streptavidin pulldown of nascent proteins. Immunoblot for newly synthesized proINS and tubulin control. (**I**) Representative immunoblots and quantification for ER stress markers with tunicamycin (TN)-treated control. Means ± SE for n = 3–4 independent experiments. *, P < 0.05 by unpaired or paired t-test.

### Ribosome profiling identifies broad impact of sustained high glucose on gene-specific mRNA translation

Incubation of MIN6 cells or islets in high glucose media supplemented with high concentrations of the saturated fatty acid, palmitate induces ER stress and has profound effects on mRNA translation (17, 18). To determine the genome-wide effects of sustained high glucose alone on β-cell mRNA translation in the absence of ER stress, we compared the translatome of MIN6 cells in low vs. high glucose by ribosome profiling. This RNA-sequencing method is based on the principle that more efficiently translated mRNAs are associated with more ribosomes and therefore generate more ribosome protected footprints (RPFs) upon nuclease digestion. Both RPF and total RNA libraries were sequenced from cells following treatment for 24 hours with 25 mM vs. 5.5 mM glucose (Figure 3A). As observed in other ribosome profiling studies of mammalian cells (23), peak RPF fragment sizes were 30-35 base pairs, RPFs were enriched for open reading frames of genes compared to mRNAs, and RPF sequences showed triplet periodicity (Figure 3B-D).

**Figure 3.**
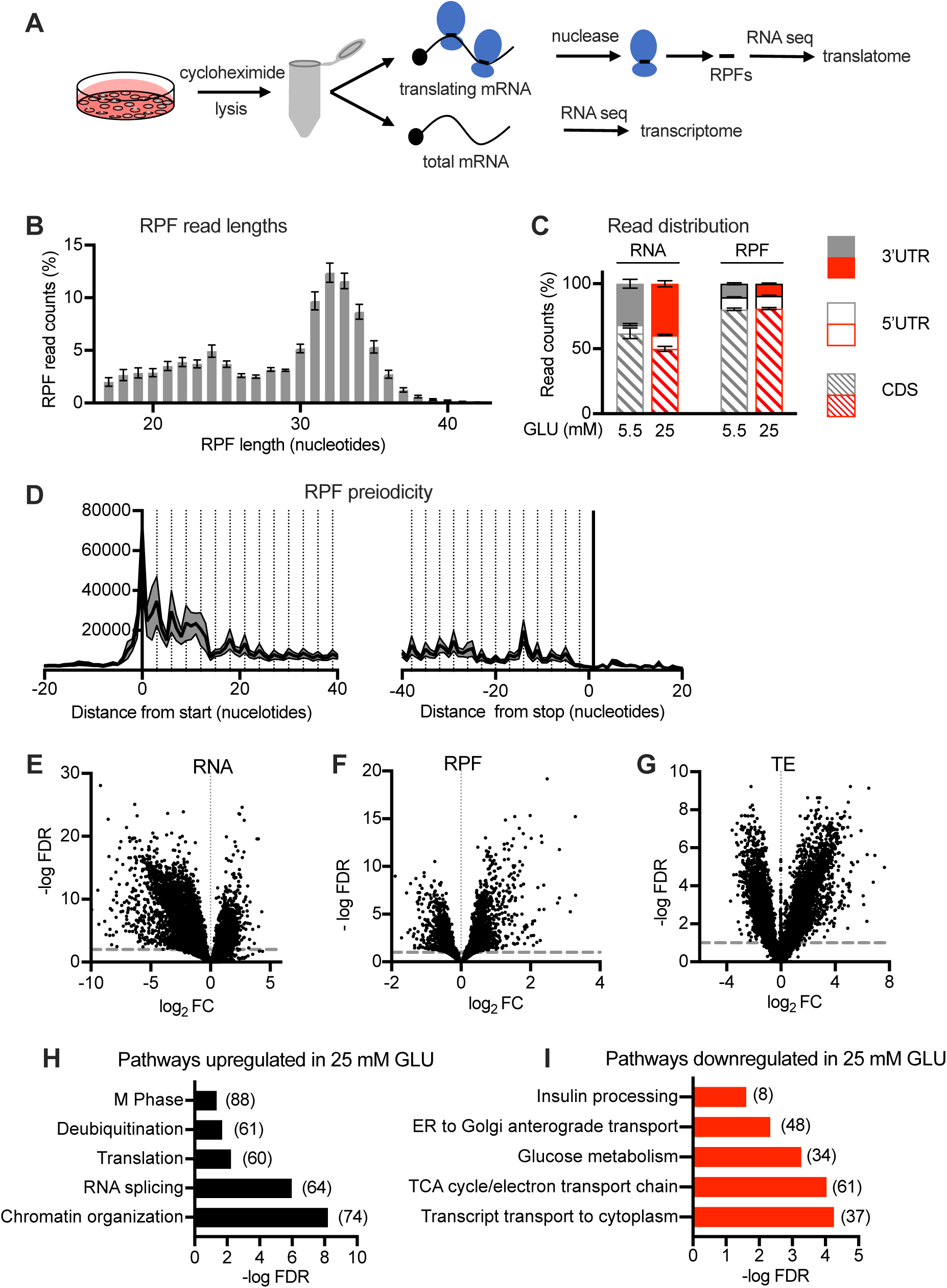
Sustained high glucose treatment has genome-wide impact on translation. MIN6 cells incubated in media containing 5.5 mM (gray) or 25 mM (red) GLU for 24 hours analyzed by ribosome profiling. (**A**) Workflow for RNA sequence analysis of ribosome protected footprints (RPFs, translatome) and total RNA (transcriptome). (**B**) RPF read lengths. (**C**) Distribution of reads to coding sequence (CDS, hatched), 5’UTR (open), 3’UTR (solid) for RNAs and RPFs. (**D**) Triplet periodicity of RPFs near CDS start and stop. (**E, F, G**) Volcano plots of-log FDR vs. log_2_ fold change (FC), calculated for 25 mM vs. 5.5 mM GLU for RNA (**E,** dotted line FDR = 0.01), RPF (**F,** dotted line FDR = 0.1) and translation efficiency (TE = RPF/RNA, **G,** dotted line FDR = 0.1). (**H, I**) Representative Reactome gene sets over-represented at a significance threshold FDR < 0.05 as upregulated (**H**) and downregulated (**I**) by 25 vs. 5.5 mM glucose. n = 8 independent samples/condition.

Sustained high glucose had substantial impact on the transcriptome (Figure 3E, 7117 genes met FDR < 0.01), consistent with prior studies (16). We observed greater representation of 3’UTR reads in the transcriptome under high glucose conditions (Figure 3C), suggesting that transcriptionally upregulated genes may have longer 3’UTR sequences. There were also many changes in the translatome that met FDR < 0.1 (Figure 3F, 2728 genes met FDR < 0.1). To identify genes for which glucose treatment specifically altered mRNA translation regulation, we calculated translation efficiency (TE) as the ratio of normalized RPF reads to normalized total mRNA reads per gene (Figure 3G). This measure accounts for changes in transcription and enables identification of genes for which changes in translation do not simply parallel changes in mRNA abundance. Using FDR < 0.1, we found that sustained high glucose up-regulated the TE of 3393 genes and down-regulated the TE of 3382 genes. Among genes for which TE was up-regulated by chronic high glucose, pathways related to chromatin organization, RNA splicing, translation, deubiquitination, and M phase of the cell cycle were over-represented (Figure 3H). Among the genes for which TE was down-regulated by chronic high glucose, pathways important for β-cell function including insulin processing, ER-to Golgi transport, glucose metabolism, and TCA cycle were over-represented (Figure 3I). Thus, independent of ER stress or change in global protein synthesis rates, sustained high glucose had genome-wide effects on translation. Moreover, in addition to insulin, sustained high glucose down-regulated translation of other genes required for metabolism-coupled insulin secretion.

### Nascent proteomics as an independent measure of glucose-altered translation

Our ribosome profiling analysis in MIN6 cells identified a large number of genes for which translation was affected by sustained high glucose based on the ratio of RPFs to total mRNA. To corroborate and filter these results, we used an orthogonal method of translation analysis in which nascent peptides were pulse labeled with the methionine analogue azidohomoalanine (AHA), click biotin conjugated, and enriched by streptavidin pulldown up front of mass spectrometry-based proteomics (Figure 4A). Principle component analysis revealed distinct patterns of nascent protein synthesis under the different glucose treatment conditions (Figure 4B). Sustained high glucose significantly affected new peptide synthesis for many genes (Figure 4C), and this correlated well with abundance of RPF sequences in ribosome profiling (Figure 4D).

**Figure 4.**
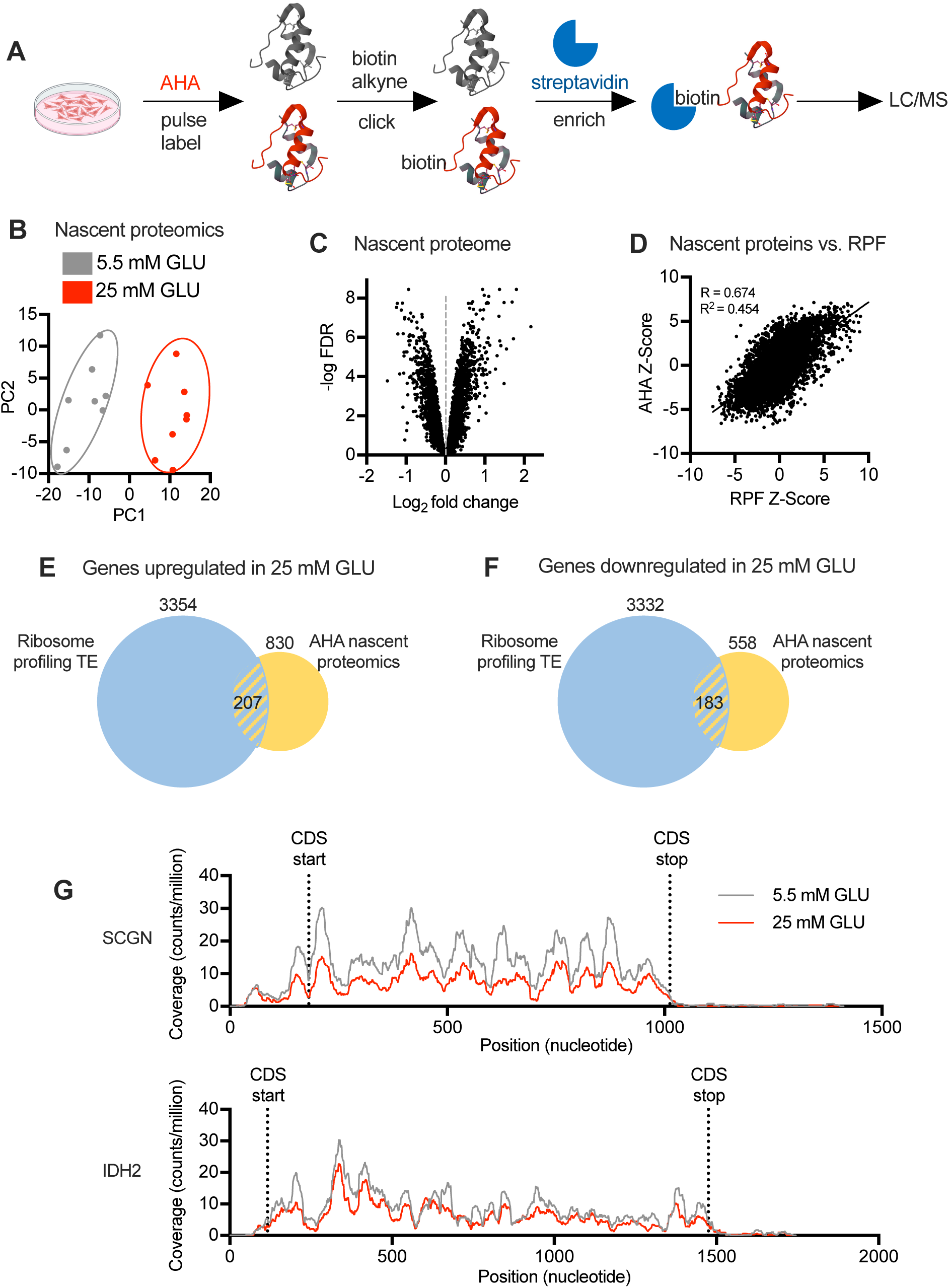
Sustained high glucose treatment has genome-wide impact on nascent proteome. MIN6 cells incubated for 24 hours in media containing 5.5 mM (gray) or 25 mM (red) GLU were analyzed by nascent proteomics. (**A**) Workflow. (**B**) Surrogate variable PCA analysis. (**C**) Volcano plots of-log FDR v. log_2_FC, calculated for 25 vs. 5.5 mM GLU. (**D**) Correlation analysis of nascent proteomics Z-scores vs. RPF z-scores. (**E, F**) Overlap of proteins upregulated (**E**) or downregulated (**F**) by 25 mM vs. 5 mM GLU in both TE and nascent proteomics datasets. –log FDR > 1 and log_2_FC > 20%. (**G**) Representative RPF gene coverage plots for SCGN and IDH2, showing no evidence for new upstream open reading frames or pausing in 25 mM GLU. n = 8 independent samples per condition. CDS, coding sequence.

To identify high confidence glucose-driven translation changes, we examined the overlap between nascent proteomics and ribosome profiling TE, focusing on proteins for which changes in both analyses met FDR < 0.1 and 20% log_2_ fold change. Based on these criteria, 207 proteins were upregulated by sustained high glucose (Figure 4E). However, this group did not contain “disallowed” genes, whose upregulation is known to lead to impaired insulin secretion (24). On the other hand, proteins known to function in insulin production and GSIS were found among the 183 downregulated proteins (Figures 4F). Given that sustained high glucose down-regulates GSIS and insulin translation, we focused on down-regulated proteins known to function in insulin production and metabolism-coupled insulin secretion (Table 1). SCGN (secretagogin) is an EF-hand Ca^2+-^ binding protein that regulates F-actin dynamics, focal adhesion remodeling, and second phase insulin secretion in β-cells, and its knockdown impairs GSIS (25–27). VPS41, a component of the homotypic function and vacuole protein sorting complex, and SLC30A8, which transports zinc into insulin granules, are both required for optimal GSIS (28, 29). SLC2A2, the plasma membrane glucose transporter in rodent β-cells, and IDH2, which functions in reductive flux of glutamine to citrate in the mitochondria, are critical for metabolism-coupled insulin secretion (30–32). While PFKFB3 and IGF2 were not detected in our nascent proteomics, they were significantly downregulated by high glucose in ribosome profiling and were included in further analyses given established roles in potentiating insulin secretion (33, 34). Prior studies show that knockdown or knockout of each of these genes diminishes GSIS with more modest or no effect on basal secretion. In our ribosome profiling analysis, RPFs for these genes were similarly distributed over the 5’ untranslated and coding regions in high and low glucose samples, without evidence for new upstream open reading frames or pausing in sustained high glucose (Figure 4G, representative traces for SCGN and IDH2).

**Table 1.** Insulin synthesis/secretion genes translationally downregulated by sustained high glucose*.

|  |  |  |  |  |  |
| --- | --- | --- | --- | --- | --- |
| Gene symbol | Role | Ribosome profiling TE |  | Nascent Proteomics |  |
|  |  | log2FC | -logFDR | log2FC | -logFDR |
| Idh2 | Insulin secretion through reductive TCA flux | -1.99 | 3.61 | -0.54 | 6.22 |
| Igf2 | Autocrine signaling promotes insulin secretion | -1.34 | 2.29 | nd | nd |
| Ins1 | Insulin gene | -0.77 | 1.14 | -0.72 | 3.61 |
| Ins2 | Insulin gene | -1.45 | 2.29 | # | # |
| Pfkfb3 | Potentiates GSIS through regulation of glucokinase | -2.43 | 4.54 | nd | nd |
| Scgn | Ca++ and insulin binding protein that enhances GSIS | -0.99 | 1.79 | -0.85 | 4.86 |
| Slc2a2 | Transports glucose for metabolism-coupled insulin secretion | -1.03 | 2.67 | -0.45 | 1.44 |
| Slc30a8 | Transports Zn++ into insulin secretory granules | -1.16 | 3.20 | -0.36 | 2.90 |
| Vps41 | Endolysosomal protein in insulin secretory granule formation | -1.26 | 3.05 | -0.41 | 4.05 |
| * Sustained high glucose decreases translation efficiency and nascent peptides by 20% or more with FDR < 0.1. nd, not detected, # detection limited by peptide overlap with Ins1 |  |  |  |  |  |

### Validation of translational changes and impact on steady state protein abundance

To confirm these chronic glucose-induced translation changes, we first quantified the ratio of mRNA associated with actively translating polysomes relative to total mRNA as a measure of TE in MIN6 cells incubated in 25 vs. 5.5 mM glucose for 24 hours. Sustained high glucose decreased TE of *Ins1*, *Ins2*, *Scgn*, *Slc2a2*, *Pfkfb3, Slc30a8, Vps41*, *Idh2*, and *Igf2*, (Figure 5A). *Actb* and *Tub1* were unchanged by high glucose, consistent with lack of change in nascent protein. To determine whether changes in TE impacted protein levels, we quantified steady-state protein abundance in lysates of MIN6 cells treated with 25 vs. 5.5 mM glucose. Sustained high glucose significantly decreased cellular content of SCGN, SLC2A2, PFKFB3, SLC30A8, VPS41, IDH2, and IGF2 (Figure 5B).

**Figure 5.**
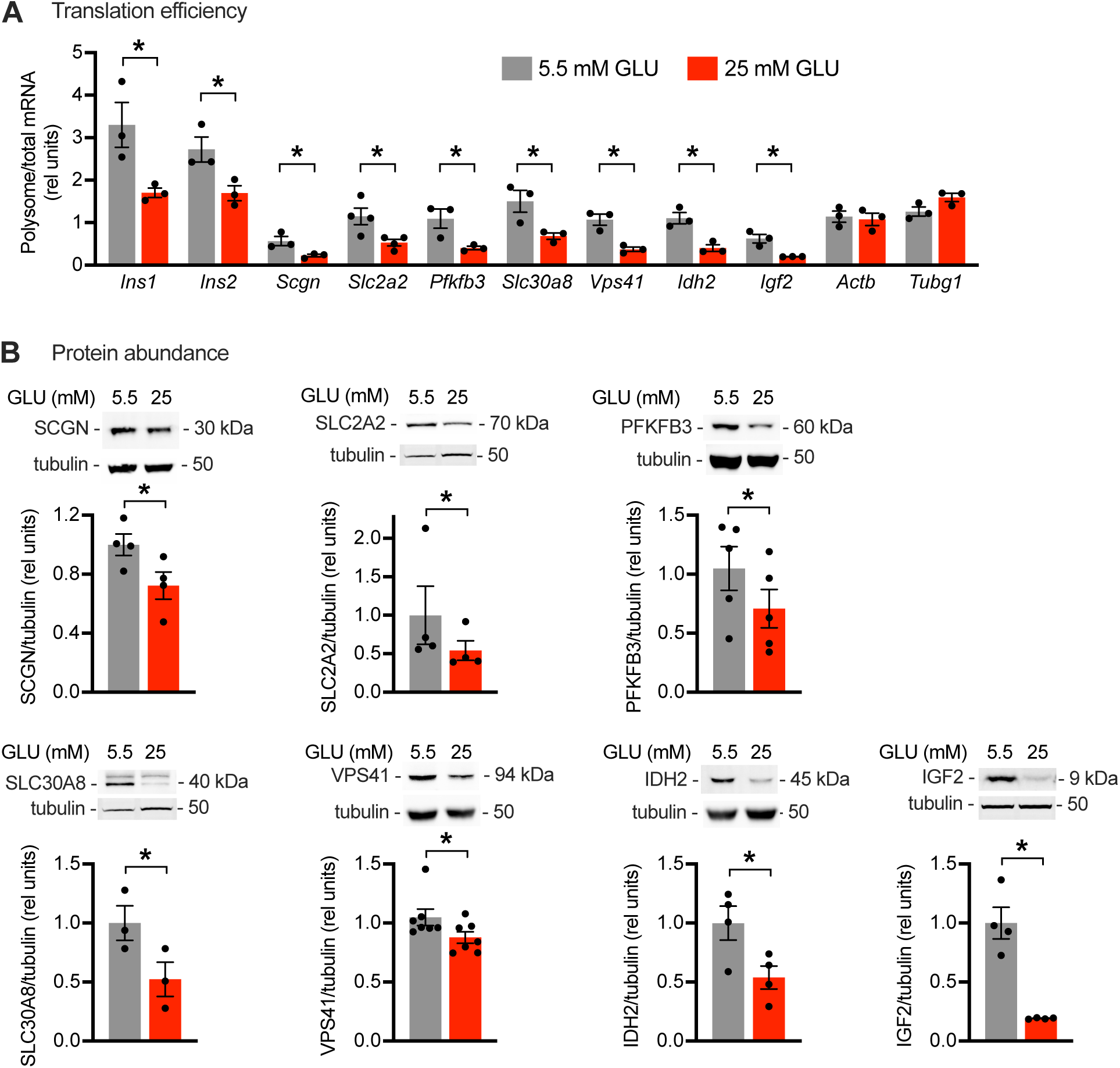
Translational regulation by sustained high glucose impacts protein abundance in MIN6 cells. MIN6 cells incubated for 24 h in media containing 5.5 mM (gray) or 25 mM (red) GLU. (**A**) qPCR quantification of polysome and total RNA for *Ins1, Ins2, Scgn, Slc2a2, Pfkfb3, Slc30a8, Vps41, Idh2, and Igf2,* relative to 18S rRNA. *Actb* and *Tubg1* as controls. Means ± SE for n = 3–4 independent experiments. *, P < 0.01 by unpaired t-test. (**B**) Immunoblot of cell lysates for SCGN, SLC2A2, PFKFB3, SLC30A8, VPS41, IDH2, and IGF2. Tubulin loading control. Representative blots with quantification of means ± SE for n = 3–7 independent experiments. *, P < 0.05 by paired t-test.

Although MIN6 cells were an important tool for technically challenging high-throughput discovery studies, these cells replicate rapidly, grow dispersed in cell culture, and lack the complex cellular make-up and architecture of islets. We next validated our findings in isolated rat islets using the conditions established above that impair glucose stimulated insulin translation and secretion (Figure 1). Given the large amount of tissue needed to collect actively translating polysomes and limited number of islets, we quantified ribosome-associated (rather than polysome) mRNA relative to total mRNA as a measure of TE. Sustained high glucose decreased TE for *Ins1*, *Ins2*, *Scgn*, *Slc2a2*, *Pfkfb3, Slc30a8, Vps41, Idh2,* and *Igf2*, without affecting TE of *Actb* and *Tubg1* (Figure 6A). As observed in MIN6 cells, this led to significantly decreased steady-state protein abundance for SCGN, SLC2A2, PFKFB3, VPS41, IDH2, and IGF2 and a trend for decrease in SLC30A8 that did not reach significance (Figure 6B). Collectively, our results in primary islets and in MIN6 cells confirm findings from ribosome profiling and provide evidence that the translational regulation we uncovered has a meaningful impact on the β-cell proteome.

**Figure 6.**
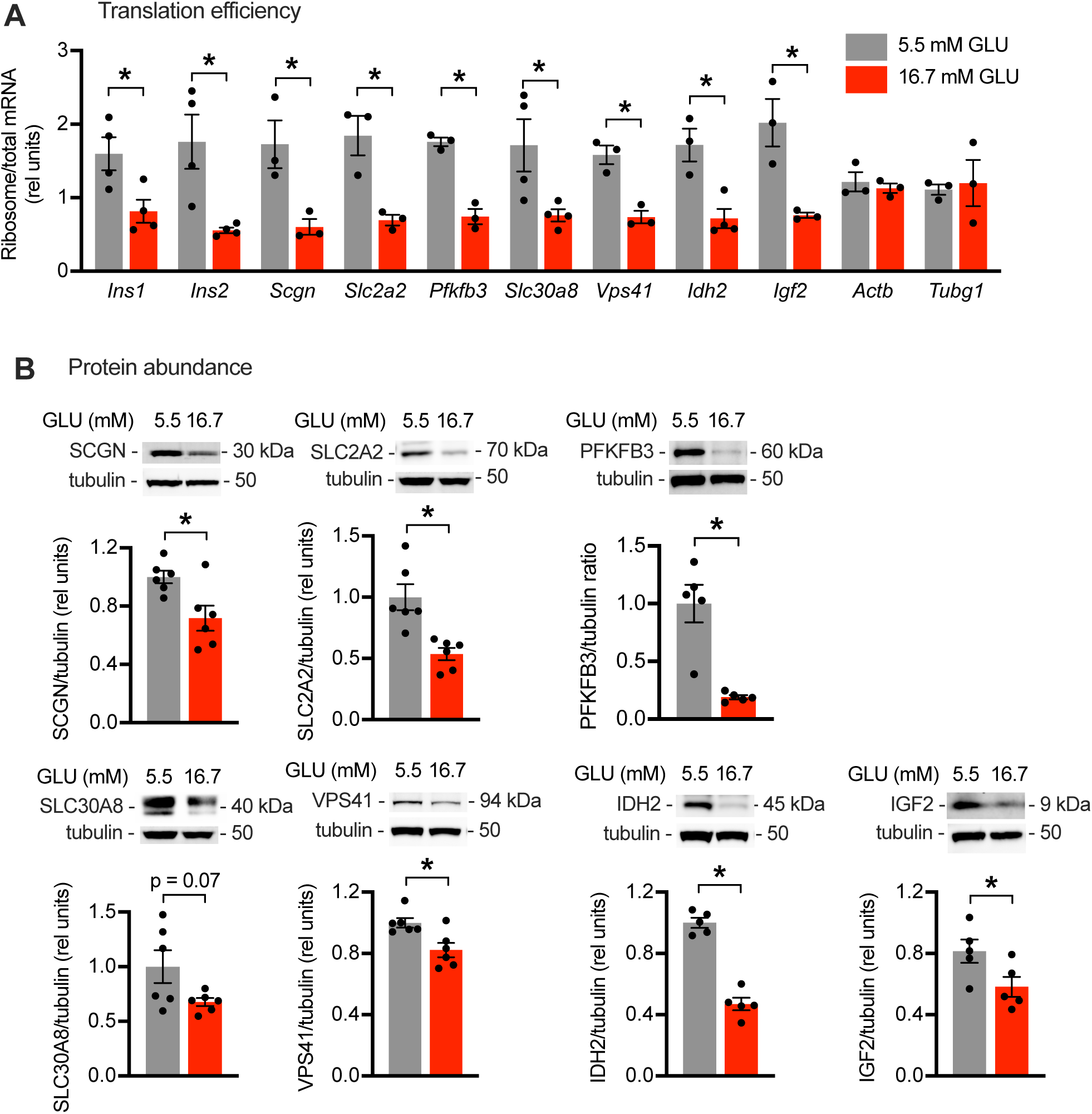
Translational regulation by sustained high glucose impacts protein abundance in rat islets. Primary rat islets incubated for 4 days in media containing 5.5 mM (gray) or 16.7 (red) mM GLU. (**A**) qPCR quantification of ribosome-associated and total RNA for *Ins1, Ins2, Scgn, Slc2a2, Pfkfb3, Slc30a8, Vps41, Idh2, and Igf2,* relative to 18S rRNA. *Actb* and *Tubg1* as controls. Means ± SE for n = 3–4 rats. *, P < 0.05 by unpaired t-test. (**B**) Representative immunoblots of islet lysates for SCGN, SLC2A2, PFKFB3, SLC30A8, VPS41, IDH2, and IGF2. Tubulin loading control. Quantification of means ± SE for n = 5–6 rats. *, P < 0.05 by paired t-test.

To investigate the clinical relevance of the observed glucose effects on translation, we analyzed TE and protein abundance following incubation of cadaveric human islets in 20 vs. 5.5 mM glucose (Figure 7A). Exposure of human islets to high glucose for 2 days was sufficient to increase basal insulin secretion and decrease stimulation index (Figure 7B), consistent with previous reports (16). Although sustained high glucose decreased insulin content of human islets (Figure 7C), GSIS was significantly decreased when normalized for islet insulin (Figure 7D). TE, as assessed by ribosome-associated/total mRNA was decreased for *INS, SCGN, PFKFB3, and VPS41* (Figure 7E). TE for *SLC30A8* trended down but did not reach statistical significance. TE for *IDH2* and *IGF2* were unchanged. Although translation of *SLC2A2* was unchanged, TE for *SLC2A1*, the main plasma membrane glucose transporter in human islets (35), was decreased. Consistent with findings in rodent islets, decreased TE led to decreased steady-state protein abundance for INS, SCGN, SLC2A1, PFKFB3, and SLC30A8 (Figure 7F).

**Figure 7.**
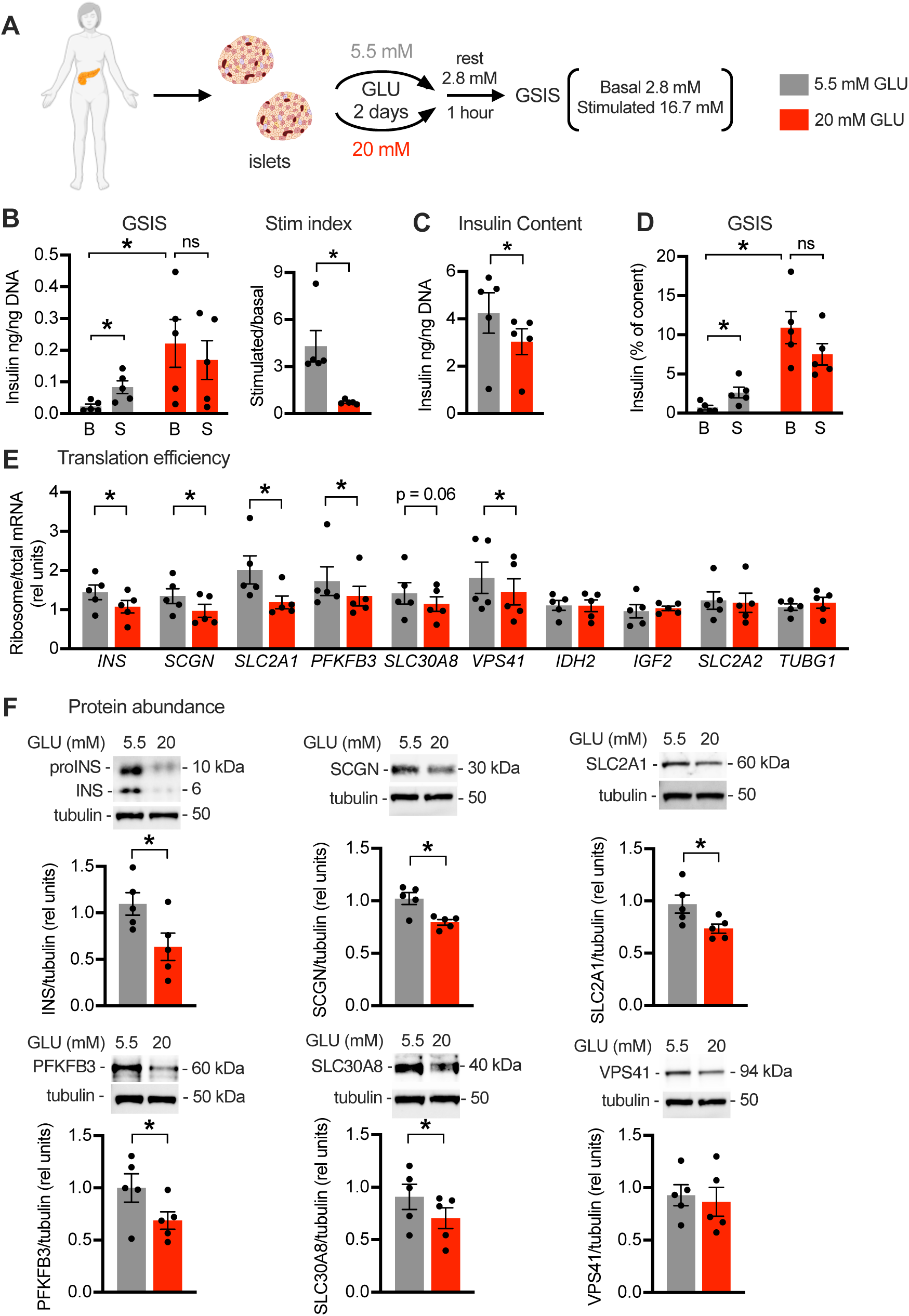
Translational regulation by sustained high glucose impacts protein abundance in human islets. Human cadaveric islets were cultured for 2 days in media containing 5.5 mM (gray) or 20 mM (red) GLU. (**A**) Following 1 hour rest in 2.8 mM GLU, GSIS quantified at 2.8 mM (B) and 16.7 mM (S) GLU. (**B**) GSIS normalized by DNA with stim index quantified as stimulatory/basal secretion. (**C**) Insulin content normalized to DNA. (**D**) GSIS normalized to cellular insulin content. (**E**) qPCR quantification of ribosome-associated and total RNA for *INS, SCGN, SLC2A1, PFKFB3, SLC30A8, VPS41, IDH2, IGF2, and SLC2A2,* relative to 18S rRNA. *TUBG1* as control. (**F**) Representative immunoblots of islet lysates for INS, SCGN, SLC2A1, PFKFB3, SLC30A8, and VPS41. Tubulin as control. Quantification of means ± SE for n = 5 donors. *, P < 0.05 by paired t-test.

To extend our findings to an *in vivo* model of hyperglycemia, we performed partial (90%) pancreatectomy (PX) or sham surgery in adult male rats (Figure 8A). Despite partial regeneration during the initial weeks of recovery, PX animals have sustained mild hyperglycemia and show selective loss of glucose-stimulated insulin secretion at 10 weeks post-surgery (36, 37). As expected, PX rats had modest, but significantly elevated, fed blood glucose compared to sham animals (Figure 8B). Islets isolated 10-weeks after PX demonstrated decreased TE for highly expressed genes including *Ins1, Ins2, Scgn, Slc2a2,* and *Slc30a8* compared to sham with no effect on TE of *Tubg1* control (Figure 8C). For genes expressed at lower levels (*Vps41, Idh2, Pfkfb3,* and *Igf2*), recovery of mRNAs was insufficient to quantify TE. Steady-state protein levels were decreased for INS, SCGN, SLC2A2, and SLC30A8 (Figure 8D). Thus, sustained exposure to high glucose in a pathophysiologically relevant setting suppressed translation of key mRNAs required for metabolism-coupled insulin secretion and abundance of their encoded proteins.

**Figure 8.**
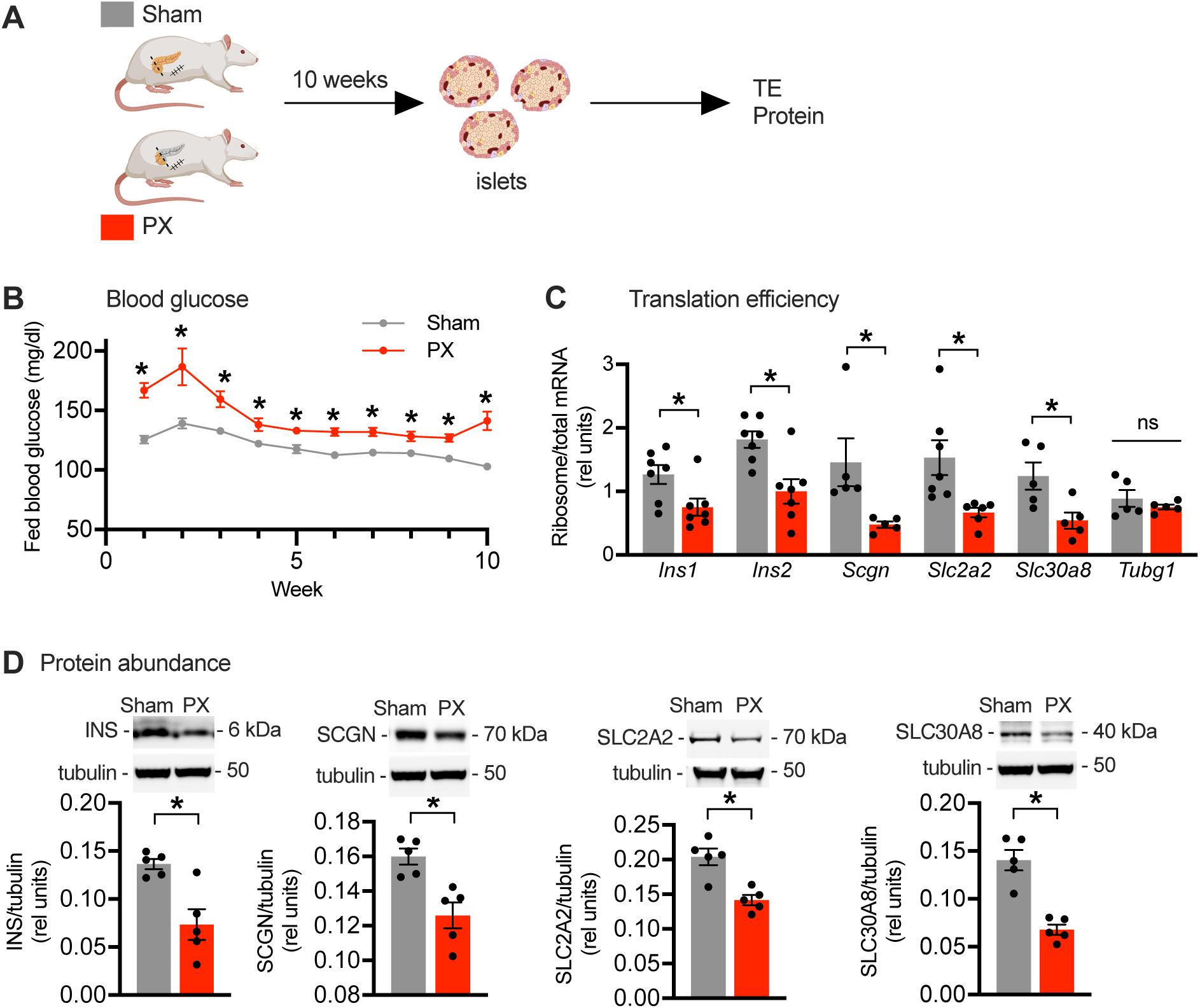
Translational regulation by hyperglycemia impacts protein abundance in partial pancreatectomy model of hyperglycemia. (**A**) Islets isolated from Sprague-Dawley rats 10 weeks after sham (gray) or 90% pancreatectomy (PX, red) surgery. (**B**) Fed blood glucose. Means ± SE for n = 21 sham rats; n = 29 PX rats. *, P < 0.05 by unpaired t-test. (**C**) qPCR quantification of ribosome-associated and total islet RNA for *Ins1, Ins2, Scgn, Slc2a2, and Slc30a8,* relative to 18S rRNA*. Tubg1 as control*. Means ± SE for n = 5–7 samples, each pooled from 2-3 rats. *, P < 0.05 by unpaired t-test. ns, not significant. (**D**) Representative immunoblots of islet lysates for INS, SCGN, SLC2A2, and SLC30A8 with tubulin as control. Quantification of means ± SE for n = 5 samples, each pooled from 2-3 rats. *, P < 0.05 by unpaired t-test.

## Discussion

In this study, we demonstrate that sustained high glucose selectively impairs mRNA translation of genes that serve critical roles at almost every step of glucose metabolism-coupled insulin secretion in pancreatic β-cells. These nutrient-induced translation changes are coincident with impaired GSIS following prolonged exposure of cultured insulinoma cells or isolated islets *ex vivo* to high glucose and in the setting of 10 weeks of systemic hyperglycemia induced by partial pancreatectomy. Our results show that programmatic dysregulation of β-cell mRNA translation is a manifestation of glucose toxicity prior to the onset of ER stress or impairment of global translation. Translational downregulation decreases steady state levels of these proteins, which serve important roles in GSIS and optimal β-cell function.

β-cells leverage structural and functional specializations for coupling glucose metabolism to robust insulin peptide production and secretion. The insulin mRNA is highly abundant, and ER and Golgi are extensive in β-cells (38, 39). Moreover, translation of insulin and genes involved in insulin processing and secretory granule biogenesis is rapidly and coordinately upregulated when glucose is acutely increased from basal to stimulatory concentrations (8, 9). Recently, high throughput studies have shown that acute exposure of β-cells to high glucose selectively upregulates translation of mRNAs encoding proteins related to insulin processing, exocytosis, and glucose metabolism, providing evidence that translation of functionally related proteins is coordinately regulated for optimal β-cell function under physiological conditions (10, 11). Our findings establish that concerted β-cell translational regulation occurs in the pathophysiological setting of chronic high glucose exposure. We show that sustained high glucose conditions that impair GSIS are associated with translational downregulation of mRNAs required for metabolism-coupled insulin secretion. Moreover, these translational changes result in decreased protein abundance, which likely contributes to decreased secretory function.

Consistent with prior reports (14, 19, 40), we found that exposure of rat and human islets to sustained high glucose decreased stimulation index but increased basal secretion. Although basal secretion and secretion following acute stimulation with glucose are supported by common components of the cellular machinery for secretion, regulation of these processes is distinct. Basal secretion is controlled by reactive oxygen species, redox regulation, and s-acylation-mediated trafficking (41), whereas GSIS is regulated by glucose metabolism, anaplerotic metabolism, the pentose monophosphate shunt, free fatty acids, the phosphoenolpyruvate cycle, and paracrine signals (42). These differences may underlie the divergent impacts of sustained high glucose on basal secretion (increased) and stimulation index (decreased). In both islets and in MIN6 cells, sustained high glucose decreased stimulation index. Lack of increase in basal insulin secretion in the MIN6 β-cell model may reflect the more modest capacity of these cells for insulin synthesis and the substantially decreased insulin content following exposure to sustained high glucose. In MIN6 and in islets, we used glucose concentrations previously demonstrated to impair GSIS (14, 16, 43). The different concentrations and durations of glucose needed to impair GSIS in insulinoma cells and islets from different species reflects intrinsic differences among these models.

β-cells synthesize up to a million proinsulin protein molecules per minute (44), creating a challenge for proper folding and processing of nascent proteins in the ER. Not surprisingly, prolonged exposure to high glucose can lead to ER-stress that activates the unfolded protein response to decrease total mRNA translation (45). Our experimental design incorporated *ex vivo* treatment of islets and MIN6 cells with glucose at concentrations and for durations that did not increase ER stress markers in order to model early nutrient-induced changes. Consistent with the lack of engagement of the PERK-eIF2 alpha arm of ER-stress, total mRNA translation was unchanged under conditions in which we observed that translation of mRNAs involved in glucose-coupled insulin secretion was suppressed (46). It is also not surprising that ATF4 and JUND, proteins for which translation is increased under glucolipotoxic conditions that induce ER stress, were not upregulated in our study (17, 18). Our results indicate that programmatic alterations in translation of specific mRNAs occurs prior to ER stress during the progression of β-cell dysfunction.

Ribosome profiling has emerged as a powerful method for assessing mRNA translation in species ranging from yeast to human (47). However, TE calculated as the ratio of RPFs to RNA is an indirect measure of translation that could be confounded by translation pausing. Our strategy to also use nascent proteomics provided an approach to filter ribosome profiling results for translation changes that resulted specifically in synthesis of new proteins. As expected, only a fraction of newly synthesized proteins reflected altered TE, since nascent proteomics does not filter out changes resulting from increased mRNA abundance (48). We validated findings from MIN6 cell discovery studies in primary rat islets in which we demonstrated significant decreases in protein abundance for translationally down-regulated genes. In the rat partial pancreatectomy model, sustained mild hyperglycemia led to decreases in translation and protein abundance for the majority of these genes for which mRNA is highly abundant. Inability to quantify less abundant RNAs was likely a consequence of our experimental design to analyze islets immediately upon isolation without overnight recovery of the islets in order to capture the impact of *in vivo* glycemia. Nonetheless, the partial pancreatectomy model provides important evidence that pathophysiological exposure to sustained high glucose over time produces similar dysregulation of islet mRNA translation as observed following *ex vivo* incubation of isolated islets.

Our human islet analyses largely phenocopied observations in rodent studies with several notable exceptions. First, TE for SLC2A1, but not SLC2A2, was significantly decreased in human islets, an observation that likely relates to species differences in glucose transporters (GLUT2/*Slc2a2* in rodent and GLUT1/*SLC2A1* in human β-cells) (35). Second, absence of change in *IDH2* and *IGF2* translation in human islets may reflect species differences in regulation of insulin secretion, as roles for these proteins in GSIS has been best characterized in rodents (31, 34). Overall, the magnitude of effect on mRNA translation in human islets was smaller than effects in MIN6 cells and rodent islets. This could relate to technical aspects of collection and handling of cadaveric tissue. Nonetheless, recapitulation of translational dysregulation by sustained high glucose in human islets supports the disease relevance of our findings.

mRNA-specific translational regulation is often driven by regulatory sequences that lie within the 5’ or 3’ UTRs of target RNAs and serve to increase or inhibit translation initiation (49). Neither mFold nor RNAfold, thermodynamic algorithms that maximize Watson-Crick base pairs and nearest-neighbor parameters, predicted common secondary structural elements in the UTRs of the mRNAs that were translationally regulated by sustained high glucose. Moreover, we did not find common RNA binding protein (RNAbp) motifs in the 5’ or 3’ UTRs of these mRNAs using the MEME-suite prediction algorithm. Because *in silico* analysis of RNA primary sequence motifs and structure have limited sensitivity and specificity (50), thorough evaluation of the contributions of UTRs to translational regulation of specific mRNAs by sustained high glucose will require in-depth experimental analyses that are beyond the scope of the present study.

Regulatory steps following transcription play an important role in determining gene expression, and simultaneous RNA sequencing and proteomic analyses combined with metabolic labeling of macromolecules provides evidence that mRNA-specific translation rates are a major determinant of the cellular proteome (51). Moreover, the development of high throughput tools for discovery of coordinated mRNA-specific translation has advanced our understanding of how environmental cues shape gene expression. Our study provides insights into nutrient-driven translational regulation that alters the abundance of proteins important for insulin secretion in settings of β-cell dysfunction. Future work to identify mechanisms by which glucose availability selectively and coordinately regulates translation of these mRNAs has the potential to identify novel strategies for interrupting β-cell dysfunction early in the progression of diabetes.

## Methods

### Rodent Islets

Islets were isolated from 7-8-week-old male Sprague-Dawley rats (Taconic Biosciences) by collagenase digestion followed by density gradient centrifugation as previously described (52). Islets were hand-picked and cultured overnight in RPMI 1640 containing 11 mM glucose, 10% FCS, 100 units/ml penicillin, 100 μg/ml streptomycin at 37 °C with 5% CO_2_. For analysis of GSIS, RNA or protein, media was changed to RPMI media containing either 5.5 mM or 16.7 mM glucose for 4 days.

### Human islets

Islets from cadaveric nondiabetic donors (ages 30-50) (Supplemental Table 1) were obtained from Prodo Labs and cultured overnight in RPMI 1640 containing 5.5 mM glucose, 10% FCS, 100 units/ml penicillin, 100 μg/ml streptomycin at 37°C with 5% CO_2_. The following day, islets were incubated in media containing either 5.5 mM or 20 mM glucose for 2 days prior to analysis of GSIS, RNA or protein.

### MIN6 cells

Low passage MIN6 cells (generously provided by Dr. Jun-ichi Miyazaki) were cultured in DMEM containing 25 mM glucose, 15% FBS, 0.1 mM β-mercaptoethanol, 100 units/ml penicillin and 100 μg/ml streptomycin at 37°C with 5% CO_2_. For analyses of translation, MIN6 cells were incubated in DMEM media containing 5.5 mM or 25 mM glucose for 24 hours prior to analysis of GSIS, RNA or protein.

### GSIS

Following incubations at different glucose concentrations, 10-12 islets (∼150 μm diameter) in triplicate or MIN6 cells (10^5^/35 mm well) in duplicate were washed and incubated with Krebs Ringer Bicarbonate HEPES buffer (KRBH: 137 mM NaCl, 4.8 mM KCl, 1.2 mM KH_2_PO4, 1.2 mM MgSO_4_ 7H_2_O, 2.5 mM CaCl_2_.2H_2_O, 5mM NaHCO_3_, 16 mM HEPES, 0.1% BSA) containing 2.8 mM glucose for 1 hour. Following media change, cells were successively incubated in KRBH containing 2.8 and 16.7 (islet) or 16.8 mM (MIN6) glucose, each for 1 hour. Media was collected for insulin quantification by Ultra Sensitive Mouse Insulin ELISA (Crystal Chem). Insulin was normalized to DNA content for islets (CyQuant cell proliferation kit, Fisher Scientific C7026) and to cell number for MIN6 cells.

### RT-qPCR quantification of mRNA

Total RNA was isolated from cell lysates, sucrose density gradient fractions, or ribosome pellets using Trizol or Trizol-LS reagents (Invitrogen) and Direct-zol RNA miniprep kit (Zymo Research). RNA recovered from sucrose density gradient fractions was treated with 600 units/ml heparinase (NEB P0735S, 20 minutes, room temperature [RT]). 500 ng to 1 µg RNA was reverse transcribed using iScript cDNA synthesis kit (Biorad). RT-qPCR was performed using SsoAdvanced Universal SYBR Green Supermix (Biorad). RNA abundance was calculated according to the ΔΔCT method relative to 18S rRNA. Primers are listed in Supplemental Table 2.

### Quantification of nascent peptides using OPP

During the last 2 hours of incubations in different media, 20 µM O-propargyl-puromycin (OPP, Click Chemistry Tools) was added to the media. Cells were lysed with RIPA buffer (50 mM Tris pH 8, 150 mM NaCl, 0.5 % sodium deoxycholate, 1% NP-40, 0.1% SDS) containing cOmplete EDTA-free protease inhibitor cocktail (Sigma). Protein was quantified using bicinchoninic acid assay (BCA, Pierce). For in-gel quantification of total nascent peptides, 100-300 µg protein were used for copper-catalyzed cycloaddition reactions using the Click-iT Plus Alexa Fluor 647 Picolyl Azide Toolkit (Fisher Scientific). Proteins were separated using NuPAGE 10% Bis-Tris gels and gels were fixed with 10% acetic acid, 50% methanol. Total nascent peptides were detected by in-gel imaging at 647 µm and total protein was detected by far red epi imaging after staining with 0.1% Coomassie brilliant blue (BioRad ChemiDoc MP Imaging System). For detection of nascent insulin and tubulin, 1 mg of cell lysate was used for cycloaddition reactions containing 5% SDS, 500 µM Biotin azide, 5 mM DTT, 0.5 mM TBTA, and 5 mM CuSO_4_ (1.5 hours, RT).

Biotinylated protein was precipitated using methanol/chloroform, re-suspended in RIPA buffer, and incubated overnight, 4°C with high-capacity streptavidin agarose beads (Pierce). Beads were washed twice sequentially with RIPA, 1 M KCl, 0.1 M Na_2_CO_3_, 2 M Urea in 50 mM Hepes, RIPA and eluted with 2x Laemmli buffer. Proteins were separated using NuPAGE 10%, Bis-Tris gels and immunoblotting was used to detect nascent proteins. See Supplemental Table 3 for antibody sources and dilutions.

### Immunoblotting

Islets were lysed by sonication in buffer containing 5 mM EDTA, 7 M urea, 2 M thiourea, 100 mM sodium fluoride, 100 mM pyrophosphate, 10 mM orthovanadate, 50 mM PMSF, 1 μg/ml aprotinin (Pierce 78432), and 1% Triton. MIN6 cells were lysed in RIPA buffer containing cOmplete protease inhibitor cocktail. 10-20 µg protein was separated on 10% NuPAGE Bis-Tris gels, transferred to PVDF membranes, blocked with 5% BSA, and blotted for the indicated proteins. Alexa Fluor-coupled (Invitrogen) secondary antibodies or HRP-coupled secondary antibodies (Cell Signaling Technologies) and chemiluminescent substrates (Biorad) were used for detection with a ChemiDoc MP Imaging System (Biorad). For ER-stress controls, islets were treated with 1 µM thapsigargin for 6 hours and MIN6 cells were treated with 5µg/mL tunicamycin for 3 hours. Antibodies and dilutions are listed in Supplemental Table 3.

### Ribosome profiling

MIN6 cells incubated in media with 5.5 mM or 25 mM glucose for 24 hours were treated with 100 µg/ml of cycloheximide for 5 minutes. Ribosome profiling was performed as previously described (53) except that 1 U/20 x 10^6^ cells RNase 1 (Thermo Scientific) was used and rDNA depletion was performed using biotinylated rDNA sequences (47). Input RNA was extracted using TRIzol (Invitrogen) and Direct-zol RNA miniprep kit. RNA libraries were generated using polyA enrichment, and Kapa stranded mRNA Hyper Prep (Illumina). RPF and RNA libraries were sequenced using Illumina NS500 single-end 75 bp reads. Data analyses employed the *XPRESSyourself* pipeline (54). Briefly, trimmed reads were aligned to the genome (Ensemble release version 102) with the two-pass option that removes rRNA alignments and PCR duplicates and counts reads that map to the exons or truncated coding sequences of the longest transcripts of protein-coding genes. *XPRESSpipe* was used for quality control analyses (RPF coverage, length and periodicity) and to obtain normalized quantification of RNA, RPF counts and TE defined as ratio of RPF to RNA. Differential expression and differential TE were performed using DESeq2 (55). Pathway analysis was performed by testing over-representation of genes with differential TE in the Reactome gene sets from MSigDB using the pre-ranked CAMERA method in the limma package with the function cameraPR (56).

### Sucrose Density Gradient Fractionation

5-50% sucrose gradients were generated using a BioComp Gradient Master IP from 5% and 50% sucrose solutions in sucrose buffer (10 Mm Tris pH 7.2, 60 mM KCl, 10 mM MgCl_2_, 1 mM DTT, and 0.1 mg/ml heparin). MIN6 cells were treated with 100 µg/ml of cycloheximide for 5 minutes. Cells were lysed with ribosome profiling lysis buffer and layered onto gradients. Following centrifugation (SW41T, 222,200 x g, 3 hours, 4°C), fractions were collected using a BR-188 Density Gradient Fractionation System (Brandel). RNA was extracted from combined polysome fractions with TRIzol LS, and cleanup used Direct-zol RNA miniprep kit.

### AHA nascent proteomics

MIN6 cells were incubated in media with 5.5 mM or 25 mM glucose for 24 hours. During the last 2.5 hours, cells were changed to methionine-free media for 30 minutes, washed with PBS and then incubated in methionine-free media containing 250 µM AHA for 2 hours. Cells were collected, lysed in RIPA buffer containing cOmplete EDTA-free protease inhibitor cocktail, and proteins were quantified by BCA. 2 mg protein per condition was reduced with 15 mM DTT (1 hour, RT), alkylated with 20 mM iodoacetamide (20 minutes, dark, RT), quenched with 10 mM DTT (15 minutes, dark, RT), precipitated using methanol/chloroform, and resuspended in 50 mM HEPES, 150 mM NaCl, 2% SDS pH 7.2. Copper-catalyzed cycloaddition of biotin was performed with 1 mg of protein by addition of 100 uM TBTA, 1 mM sodium ascorbate, 1 mM copper sulfate, 100 uM biotin-alkyne (2 hours, RT). Proteins were precipitated to remove excess biotin-alkyne, re-suspended in 2% SDS, 5 mM DTT, and diluted with RIPA buffer to final SDS to < 0.5%. Samples were mixed with 10 ul of high-capacity streptavidin beads (overnight, RT) and then washed twice sequentially with RIPA, 1 M KCl, 0.1 M Na_2_CO_3_, 2 M Urea in 50 mM Hepes, RIPA, and PBS pH 7.4. Tryptic digest, TMT labeling, separation into 6 fractions, and LC-MS3 analysis was performed as described (57). MS2 spectra were searched using the COMET algorithm against a Uniprot composite database derived from the mouse proteome, exogenous sequence, known contaminants, and reverse sequences. Peptide spectral matches were filtered to a 1% FDR using the target-decoy strategy combined with linear discriminant analysis. The proteins from the 6 runs were filtered to a <1% FDR. At least 2 unique peptides were required for identification, and proteins were quantified only from peptides with a summed SN threshold of >150. Protein intensity was log_2_ transformed, missing values imputed, and data was normalized such that all samples had the same median abundance (58). We used limma to perform linear modeling and moderated t-tests, with adjustment for surrogate variable analysis as previously described (59, 60).

### Partial pancreatectomy

Six-week-old Sprague-Dawley (∼100 g) male rats underwent 90% pancreatectomy or sham surgery as previously described (36). Under anesthesia with ketamine/xylazine, pancreatic tissue was removed by gentle abrasion with cotton-tipped applicators, leaving a small remnant 1-2 mm from the common bile duct and extending to the first loop of the duodenum. For sham surgery, the pancreas was disengaged from the mesentery but not removed. Post-operatively, body weights and morning fed glucose values were measured weekly. 10 weeks following surgery, islets were isolated as above and immediately lysed for analysis of total and ribosome-associated RNA and protein expression analysis.

### Isolation of ribosome-associated mRNA from rat and human islets

Following *ex vivo* incubation of rat and human islets with low or high glucose or immediately following isolation of islets from sham and PX rats, islets were treated with 100 µg/ml of cycloheximide for 5 minutes, washed with ice-cold PBS, lysed with ribosome profiling buffer. One-third of the sample was collected for total RNA isolation and the remaining material was centrifuged through a 1M sucrose cushion to collect pelleted ribosome-associated mRNA (435,400 x g, 1 hour, 4°C). Total and ribosome-associated mRNA were isolated using TRIzol (Invitrogen) and 1-bromo-3-chloropropane (Sigma) using phase separation method.

### Statistics

For biochemical, cell biological, and physiological experiments, analyses were performed using GraphPad Prism. Data are presented as means ± SE. Statistical significance was determined by unpaired or paired 2-tailed t-tests, as described on figure legends and p value < 0.05 was considered significant. For ribosome profiling and nascent proteomics, p-values were adjusted for multiple tests and FDR < 0.1 (Benjamini-Hochberg method) was considered significant. Figures were generated using GraphPad Prism and BioRender.

### Study Approval

All procedures using animals were approved by the Joslin Institutional Animal Care and Use Committee. The Joslin Committee on Human Studies determined that studies of de-identified cadaveric human islets do not constitute human subject research.

### Data Availability

RNA-seq data will be deposited in the NCBI Gene Omnibus (GEO) database. Proteomics data will be uploaded to the ProteomeXchange Consortium via PRIDE. Accession numbers will be provided.

## Author Contributions

AC designed the study, performed the experiments, and wrote the manuscript. JHL and BS helped perform the experiments and edited the manuscript. HP and JMD performed bioinformatic analyses. SBW provided mentorship and edited the manuscript. JES designed the study, provided mentorship, and wrote the manuscript.

## Acknowledgments

We are grateful to Dr. Jun-Ichi and Dr. Satsuki Miyazaki for the generous gift of low passage MIN6 cells, and to Dr. Cristina Aguayo-Mazzucato and Dr. Gordon Weir for helpful discussions and critical reading of the manuscript. This work was supported by NIH DP1 DK119141 (JES), Beatson Foundation Award 2022-005 (JES), NIH T32 DK00760 (AC), Joslin Diabetes Research Center (2P30DK036836) Molecular Phenotyping and Genotyping Core, the Joslin Islet Isolation and Bioinformatics and Biostatistics Cores, and the Diabetes Research and Wellness Foundation (SBW).

